# *Fitm2* is required for ER homeostasis and normal function of murine liver

**DOI:** 10.1101/2022.05.24.493342

**Authors:** Laura M. Bond, Ayon Ibrahim, Zon W. Lai, Rosemary L. Walzem, Roderick T. Bronson, Olga R. Ilkayeva, Tobias C. Walther, Robert V. Farese

## Abstract

The ER-resident protein fat-inducing transcript 2 (FIT2) catalyzes acyl-CoA cleavage in vitro and is required for endoplasmic reticulum (ER) homeostasis and normal lipid storage in cells. The gene encoding FIT2 is essential for the viability of mice and worms. Whether FIT2 acts as an acyl-CoA diphosphatase in vivo and how this activity affects liver, where the protein was discovered, are unknown. Here, we report that hepatocyte-specific *Fitm2* knockout (FIT2-LKO) mice exhibited elevated acyl-CoA levels, ER stress, and signs of liver injury. FIT2-LKO mice fed a chow diet had more triglycerides in their livers than control littermates due, in part, to impaired secretion of triglyceride-rich lipoproteins and reduced capacity for fatty acid oxidation. Challenging FIT2-LKO mice with a high-fat diet to increase FIT2 acyl-CoA substrates worsened hepatic ER stress and liver injury, but unexpectedly reversed the steatosis phenotype, similar to what is observed in FIT2-deficient cells loaded with fatty acids. Our findings support the model that FIT2 acts as an acyl-CoA diphosphatase in vivo and is crucial for normal hepatocyte function and ER homeostasis in murine liver.

## Introduction

The endoplasmic reticulum (ER) is the major cellular site of lipid synthesis and production of cell surface and secreted proteins. The protein fat-inducing transcript 2 (FIT2) has emerged as an important determinant of ER homeostasis in cells. FIT1 and FIT2 genes were identified as targets of the transcription factor PPARα in murine liver (1). FIT1 and FIT2 are ER-resident proteins with six putative transmembrane segments and ∼50% amino acid sequence similarity (2). Murine FIT1 and FIT2 have different tissue expression patterns: FIT1 is expressed mainly in skeletal and cardiac muscle, and FIT2 is ubiquitously expressed, with highest levels in adipose tissue (1).

The molecular functions of FIT proteins have been somewhat enigmatic. FIT2 was initially implicated in lipid metabolism and, in particular, lipid droplet (LD) biogenesis (1,3). Overexpression of FIT2 in murine liver results in increased lipid storage in hepatocytes, and this finding was replicated in a variety of cell types (1). In cells, FIT2 was localized to the ER and found at sites of LD biogenesis (4). FIT2 binds neutral lipids and was hypothesized to partition neutral lipids into a storage pool (5). The FIT2 gene is essential in worms and mice (6,7), highlighting the importance of FIT2 function. Tissue-specific deletions of *Fitm2* in mice revealed crucial functions in adipocyte differentiation, enterocyte function, and pancreatic β-cells (7–9). In humans, homozygous *FITM2* deficiency causes deafness-dystonia (10,11), and human FIT2 is required for cancer cell fitness during exposure to interferon-gamma (IFNγ) (12). The diverse deletion phenotypes show that FIT2 is crucial for life and highlight that FIT2 deficiency manifests differently in different biological systems.

FIT2 is required for normal cellular ER homeostasis. It was identified as a putative lipid phosphate phosphatase enzyme by homology searches (13), and it has acyl-CoA diphosphatase activity in vitro, utilizing a variety of acyl-CoA substrates to generate acyl 4′-phosphopantetheine and adenosine-3′,5′-bisphosphate products (14). This enzymatic activity is critical to preserving cellular ER homeostasis, as FIT2 deficiency in mammary carcinoma cells and yeast results in ER dilation and whorls, ER stress, and reduced LD biogenesis capacity (6,14). Consistent with these data, FIT2 orthologs in yeast, *SCS3* and *YFT2*, were implicated in ER homeostasis (15,16).

In the current study, we sought to determine whether FIT2 acts as an acyl-CoA cleaving enzyme and functions in ER homeostasis in vivo by deleting *Fitm2* in in murine hepatocytes. We studied the effects of FIT2 deficiency on hepatic ER and lipid homeostasis by analyzing the phenotypes of mice fed chow or high-fat diets. Our results demonstrate the necessity of FIT2 for ER homeostasis in vivo and provide insights into its physiological functions in this tissue.

## Results

### Generation of hepatocyte-specific FIT2 knockout mice

To investigate the function of FIT2 in murine liver, we generated hepatocyte-specific FIT2 knockout mice (FIT2-LKO) using Cre-loxP technology. We crossed *Fitm2*^*loxP/loxP*^ (floxed) mice with mice expressing Cre recombinase under control of the albumin promoter, leading to deletion of exon 1 of *Fitm2* (Figure S1A). Hepatic *Fitm2* transcript and protein levels, as measured by immunoblot analysis, were reduced by 95% (Figure 1A and 1B). Mass spectrometry of liver lysates also demonstrated the loss of FIT2 protein in FIT2-LKO livers (Table S1). We confirmed that Cre recombinase expression was restricted to liver and not found in muscle or adipose tissue of FIT2-LKO mice (data not shown). Consistently, *Fitm2* mRNA levels were normal in skeletal muscle, brown adipose tissue, gonadal and inguinal white adipose tissue (Figure 1A). To determine whether hepatocyte FIT2 deletion resulted in compensation by FIT1, we measured FIT1 protein levels by mass spectrometry. FIT1 was present in skeletal muscle, but was not detected in livers of either floxed control or FIT2-LKO mice, consistent with reports that FIT1 protein is not expressed in murine liver (Table S1) (1).

**Figure 1.**
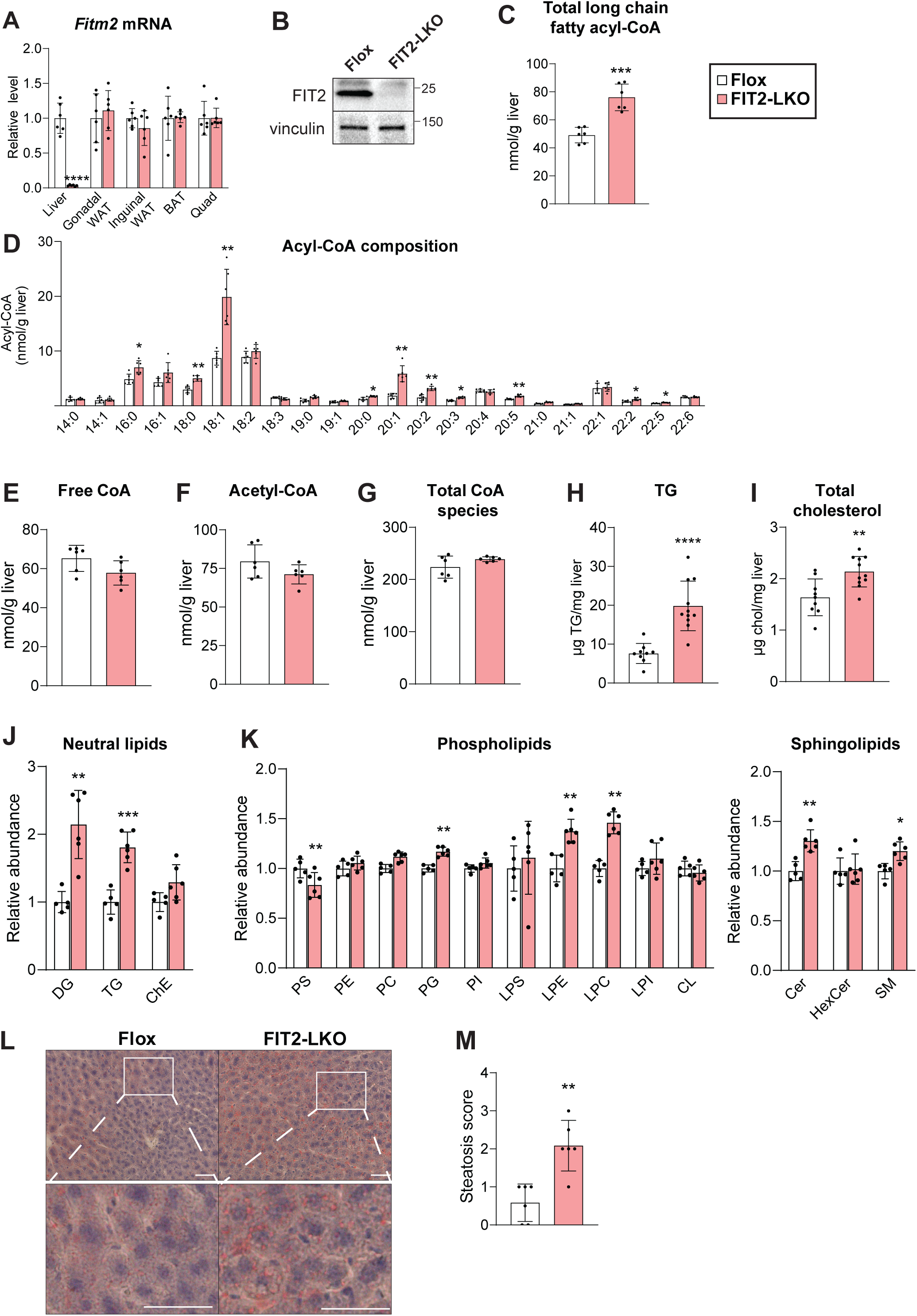
Hepatocyte-specific FIT2-LKO mice have altered hepatic lipid composition. (A) Hepatocyte-specific FIT2 knockout results in near loss of FIT2 transcript in liver (n=9-11/genotype) but does not alter FIT2 transcript levels in adipose tissue and muscle (n=6/genotype) as measured by qRT-PCR. (B) Immunoblotting indicates loss of FIT2 protein in liver tissue. (C-G) MS-based measurements of CoA and CoA derivatives in livers of Flox and FIT2-LKO mice (n=6/genotype). (C) FIT2-LKO livers contain elevated levels of total long-chain fatty acids (≥14 carbon length). (D) Amounts of individual long-chain acyl-CoA species in livers of Flox and FIT2-LKO mice. Levels of hepatic free CoA (E), acetyl-CoA (F), and total CoA-containing species (G) are not significantly altered between Flox and FIT2-LKO mice. Hepatic triglyceride levels (H) and cholesterol levels (I) are elevated in FIT2-deficient livers (n=9-11/genotype). Lipidomic analyses of levels of neutral lipids (J) and phospholipids (K) in male FIT2-LKO mice (n=6/genotype). Representative images (L) and steatosis scoring (M) of Oil Red O staining indicates more neutral lipid content in FIT2-LKO livers than in Flox livers. (n=6/genotype). Scale bar=50 µm. Data represents mean ± SD. Statistical significance for was evaluated with unpaired Student’s 2-tailed t-test for (A-K) and a Mann-Whitney U test for nonparametric data in (K). *p<0.05, **p<0.01, ***p<0.001, ****p<0.0001. TG=triglyceride; ChE=cholesterol ester; DAG=diacylglycerol; PS=phosphotidylserine; PG=phosphotidylglycerol, PE=phosphotidylethanolamine; PC=phosphotidylcholine, L=lyso, PI= phosphatidylinositol; CL=cardiolipin, Cer=ceramide; CerG1=glucosylceramide; SM=sphingomyelin.

### FIT2 deficiency alters acyl-CoAs and other hepatic lipids

To determine if FIT2 functions as an acyl-CoA diphosphatase in hepatocytes, we tested whether FIT2 deficiency leads to accumulation of acyl-CoA substrates. We measured acyl-CoA and CoA levels in control and FIT2-LKO livers (Table S2) and found that long-chain fatty acyl-CoA levels were elevated by ∼60% (Figure 1C). Specifically, we found marked increases in the levels of several species of acyl-CoAs, including the monounsaturated fatty acyl-CoAs 18:1 and 20:1, and lesser increases in several minor species (Figure 1D). In contrast, levels of free CoA, acetyl-CoA, and total CoA-containing species were normal (Figure 1E, 1F, 1G, Table S2). These data are consistent with previous findings that FIT2 hydrolyzes long-chain unsaturated fatty acyl-CoAs, but not short-chain acyl-CoA species or free CoA (14).

Because FIT2 functions in fatty acid metabolism, we also investigated changes in hepatic lipids. In contrast to reduced triglyceride (TG) storage in cells with FIT2 deficiency (1,3,14), hepatic triglyceride content was unexpectedly elevated ∼2.5-fold, and cholesterol content was increased ∼20% in both male and female mice (Figure 1H, 1I, S1B). Mass spectrometry–based lipidomic analyses of livers of control and FIT2-LKO mice corroborated the elevated triglyceride levels and revealed a twofold increase in diacylglycerol levels (Figure 1J). Levels of the major phospholipids phosphatidylcholine (PC) and phosphatidylethanolamine (PE) were not altered, but levels of several other phospholipids (e.g., lysophosphatidylcholine, lysophosphatidylethanolamine, and phosphotidylglycerol) were slightly elevated, and phosphotidylserine levels were modestly reduced (Figure 1K). Levels of ceramide and sphingomyelin were modestly elevated (Figure 1K).

We analyzed lipid deposition in histological sections with Oil Red O and H&E staining (Figure 1L, S1C). Neutral lipid deposition was low in both genotypes and agreed with minimal lipid accumulation noted under chow feeding. However, steatosis scores were greater in FIT2-LKO mice than in floxed control mice (Figure 1M). The increase in hepatic triglycerides was accompanied by elevated liver weights in both males and females (Figure S1D). Body weights were not altered between the genotypes after chow feeding (Figure S1E). Collectively, these results show that liver-specific FIT2 deficiency increases neutral lipid (i.e., TG) storage under conditions of chow feeding.

### Loss of FIT2 disrupts ER homeostasis and causes liver injury

Since FIT2 is crucial for maintaining ER homeostasis in cultured mammalian and yeast cells, we examined ER morphology and function in FIT2-LKO mice. FIT2-LKO mice exhibited modest ER stress, reflected in increased mRNA levels of transcription factors (e.g., *Xbp1*, A*tf3, Chop*) and chaperones (e.g., *BiP*) associated with the unfolded protein response in FIT2-LKO livers (Figure 2A). Phosphorylation of eIF2α was ∼three-fold greater in livers of FIT2-LKO mice than in floxed control littermates (Figure 2B). Analysis of hepatocyte ER structure by electron microscopy (EM) showed that hepatic FIT2 deficiency did not cause detectable ER dilation (Figure S2A,B). Total ER content was also apparently unaltered, as quantified by EM (Figure S2C) and proteomic analysis of ER markers (Figure S2D). Consistent with these measurements, the levels of PC and PE, major phospholipids constituents of the ER, were not altered (Figure 1K).

**Figure 2.**
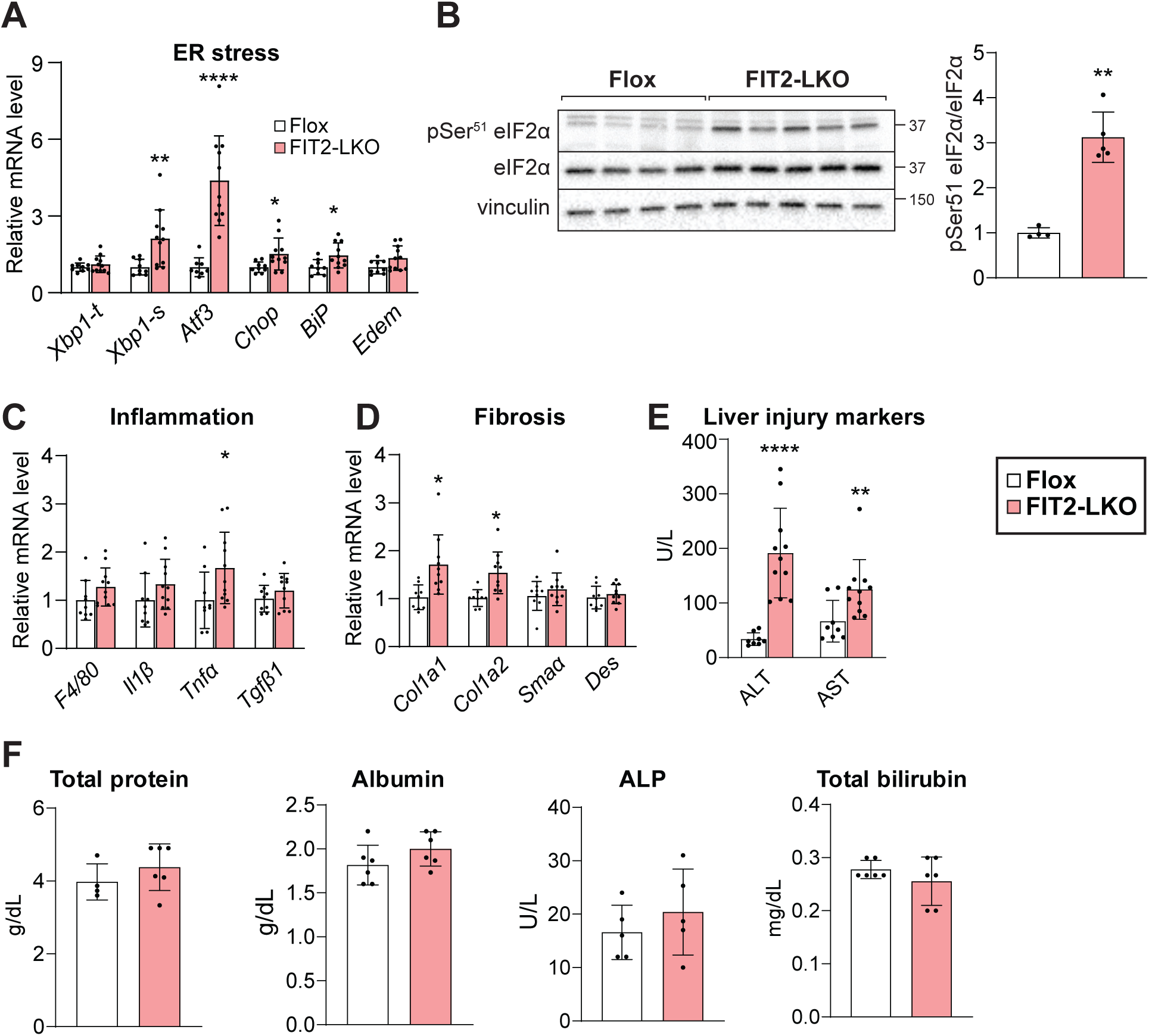
FIT2-LKO mice exhibit increased ER stress and liver injury. (A) Gene expression analyses demonstrate that FIT2-LKO mice have elevated transcript levels of chaperones and transcription factors ciated with ER stress compared to chow-fed Flox mice (n=9-11). (B) Immunoblotting indicates more phosphorylation of eIF2α in FIT2-LKO livers than in Flox littermates (n=3-4). (C) FIT2-LKO livers do not show much evidence for inflammation, as indicated by transcript levels of macrophage markers and kines (n=9-11/genotype). (D) FIT2-LKO livers do not show evidence for substantial fibrosis -11/genotype). (E) FIT2-LKO mice exhibit liver injury. Levels of circulating alanine transaminase T) and aspartate transaminase (AST) are higher in FIT2-LKO mice than in Flox controls (n=9-11). (F) ma albumin, total protein, alkaline phosphatase (ALP) and bilirubin are unaltered in FIT2-LKO mice). Data represents mean ± SD. Statistical significance for (A, C, D) was evaluated with unpaired ent’s 2-tailed t-test for parametric data and a Mann-Whitney U test for nonparametric data. Statistical ificance for (B) was evaluated with unpaired Student’s 2-tailed t-test. For (E and F), 2-way ANOVA Šidák correction was used. *p<0.05, **p<0.01, ***p<0.001, ****p<0.0001.

Because inflammation often accompanies ER stress, we assessed hepatic immune cell infiltration and cytokine production. Consistent with chow feeding eliciting minimal inflammation and negligible fibrosis, transcript levels of these markers were low in both genotypes (data not shown). However, transcript levels of macrophage markers and cytokines trended higher in FIT2-deficient livers (Figure 2C). Also, some markers of fibrosis, and transcript levels of pro- and anti-apoptotic genes were elevated (Figure 2D, S2E, S2F).

Despite minimal evidence for inflammation or apoptosis, FIT2-deficient livers displayed evidence of injury. Plasma levels of transaminases alanine transaminase and aspartate transaminase were elevated seven- and twofold, respectively (Figure 2C). Plasma markers of synthetic liver function (total protein, albumin) and cholestasis (bilirubin, alkaline phosphatase) were normal (Figure 2D).

### Impaired TG secretion and reduced fatty acid oxidation capacity contribute to TG accumulation in chow-fed FIT2-LKO mice

To elucidate the causes of hepatic TG accumulation found in chow-fed FIT2-LKO mice, we analyzed pathways that influence TG levels. We evaluated very low-density lipoprotein (VLDL) secretion, since this pathway depends on ER phospholipid and protein composition and occurs in the ER lumen (17,18), the proposed location of the catalytic residues of FIT2 (2,14). Steady state plasma levels of TG were unaltered, and levels of the primary protein component of VLDL, apolipoprotein (apo)-B, were increased in FIT2-LKO animals (Figure 3A and 3B). LDL-cholesterol levels were also similar in both genotypes (Figure 3C). However, we found that TG secretion by the liver was reduced by ∼30% in the FIT2-LKO mice (Figure 3D). In contrast, secretion of apoB, quantified by immunoblotting, was similar between genotypes (Figure 3D, Figure S3A). Proteomic analyses of the livers indicated that protein levels of apoB and the microsomal TG transfer protein (MTTP), required for lipidation of apoB, were similar among genotypes (Figure S3B). Since circulating apoB was consistently elevated and TG secretion was reduced in FIT2-LKO mice, we hypothesized that FIT2-LKO hepatocytes secrete smaller VLDL particles. To test this, we determined the size distribution of particles recovered from the *d* < 1.063 g/mL fraction of plasma. We found an increase in total percentage of particles ≤ 51 nm in diameter (88% v. 72%) and a decrease in particles ≥ 72 nm in diameter (12% v. 27%) in FIT2-LKO plasma, consistent with this hypothesis.

**Figuure 3.**
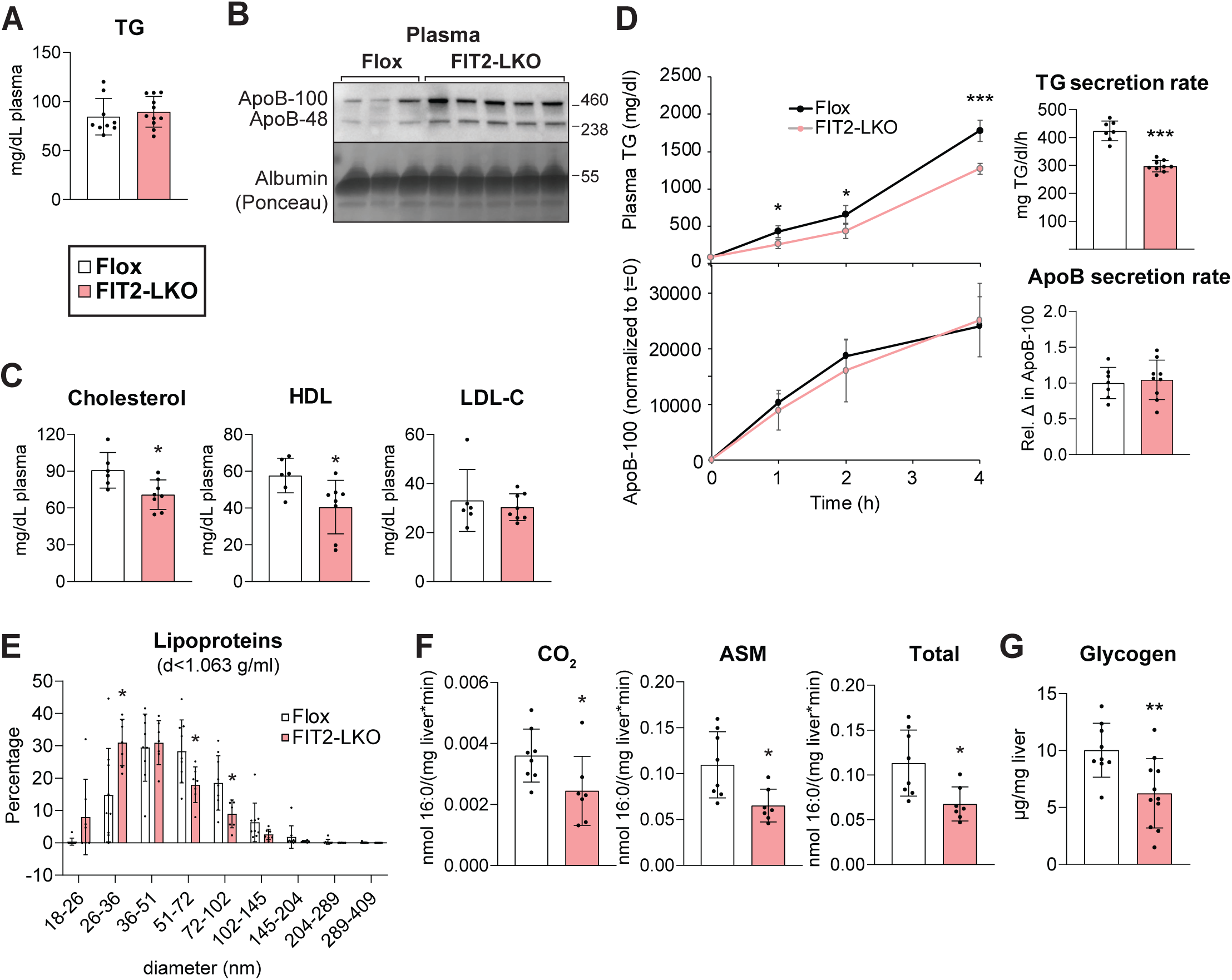
Chow-fed FIT2-LKO mice exhibit alterations in TG metabolism that contribute to steatosis. (A) Steady-state plasma TG levels are unaltered in Flox and FIT2-LKO mice (n=9-11/genotype). (B) ady-state levels of plasma apoB-100 and apoB-48 are higher in plasma of FIT2-LKO mice than in Flox ntrols. Mice were fasted 2 h before sacrifice in (A) and (B). (C) Plasma cholesterol levels were decreased the FIT2-LKO; specifically, HDL-C was reduced, but LDL-C remained the same. (D) FIT2-LKO mice hibit reduced rates of hepatic TG secretion but unaltered rates of hepatic apoB secretion. Plasma was lected before (t=0) and 1, 2, and 4 h after intravenous administration of polaxomer-407, a lipoprotein ase inhibitor. Plasma TG was assayed biochemically. Relative plasma apoB-100 protein levels were ermined by quantification of apoB-100 band intensity on immunoblots (Figure S4A). (E) Dynamic light ttering measurements indicate that lipoprotein (density <1.063) particle diameter (number distribution) reduced in FIT2-LKO mice compared to Flox mice (n=7–9/genotype) (mice were fasted 4-h before rifice). (F) FAO capacity is reduced in FIT2-LKO liver lysates. (n=5–8/genotype) (G) Hepatic glycogen els are reduced in male FIT2-LKO mice compared to Flox mice (n=9–11/genotype). Data represents an ± SD. Statistical significance for (A, C, F, and G) was evaluated with unpaired Student’s 2-tailed t-test. r (D, left), repeated-measures ANOVA was used and for (D, right), unpaired Student’s 2-tailed t-test was d. For (E), 2-way ANOVA with Šidák correction was used. *p<0.05, **p<0.01, ***p<0.001.

We also investigated whether impaired fatty acid oxidation contributes to the increased steatosis of chow-fed FIT2-LKO. Testing mitochondrial fatty acid oxidation was compelling since the deafness-dystonia syndrome reported in humans with *FITM2* mutations is reminiscent of a similar disorder, Mohr-Tranebjaerg syndrome, that is caused by defects in mitochondrial function (19,20). Liver lysates from FIT2-LKO mice had reduced capacity to produce acid-soluble metabolites and CO_2_ by oxidizing fatty acids (Figure 3F). This was not due to a reduction in transcript or protein levels of fatty acid oxidation enzymes or mitochondrial content, as assessed by oxidative phosphorylation gene expression, protein levels, and mitochondrial DNA content (Figure S3C-G). Impaired fatty acid oxidation appeared to alter fuel utilization and increased reliance on glucose oxidation in FIT2-LKO. Electron microscopy revealed a marked decrease in glycogen in hepatocytes of FIT2-LKO mice (Figure S2A), which was corroborated with a biochemical assay for hepatic glycogen (Figure 2G).

### High-fat diet worsens liver injury in FIT2-LKO mice

To further test the role of FIT2 in liver lipid and ER homeostasis, we challenged mice with a high-fat diet (HFD, 42% kcal from fat). We hypothesized that this diet would result in sustained acyl-CoA overexposure and exacerbate the phenotypes found with standard chow feeding. With respect to general parameters, FIT2-LKO mice unexpectedly gained less weight than control littermates during the 11-week feeding study (Figure S4A). The reduced weight gain was due to reduced body fat, with both gonadal and inguinal white adipose tissues showing reduced mass in FIT2-LKO mice (Figure S4B). Weekly food consumption was similar among genotypes, suggesting that increased energy expenditure led to the lower body weight phenotype (Figure S4C). We did not investigate this aspect of the phenotype further, but hepatic injury may have resulted in more energy expenditure.

In contrast to the results with a chow feeding, hepatic TG content was ∼50% less in the FIT2-LKO mice fed the HFD than in controls (Figure 4A). This was accompanied by reductions in plasma TGs and cholesterol (Figures 4B-C). The decreased hepatic lipid content was visible in H&E-stained liver tissue sections and scoring of Oil Red O staining (Figure 4D-E, S4H). Examination of the lipid deposition revealed that Flox control mice exhibited extensive centrilobular microsteatosis; in contrast, the FIT2-LKO livers exhibited predominately macrosteatosis and fat accumulation localized to the periportal zone. Consistent with the findings of reduced lipid levels, FIT2-LKO livers also showed a decrease in the expression of genes of *de novo* lipogenesis (Figure S4D). FIT2-LKO also exhibited reduced liver glycogen levels (Figure S4E), consistent with the hypothesis that they utilize carbohydrates for fuel. HFD-fed FIT2-LKO mice exhibited elevated plasma ketone bodies (Figure S4F), although they showed little to no differences in the expression of fatty acid oxidation or oxidative phosphorylation-related genes (Figure S4G).

**Figure 4.**
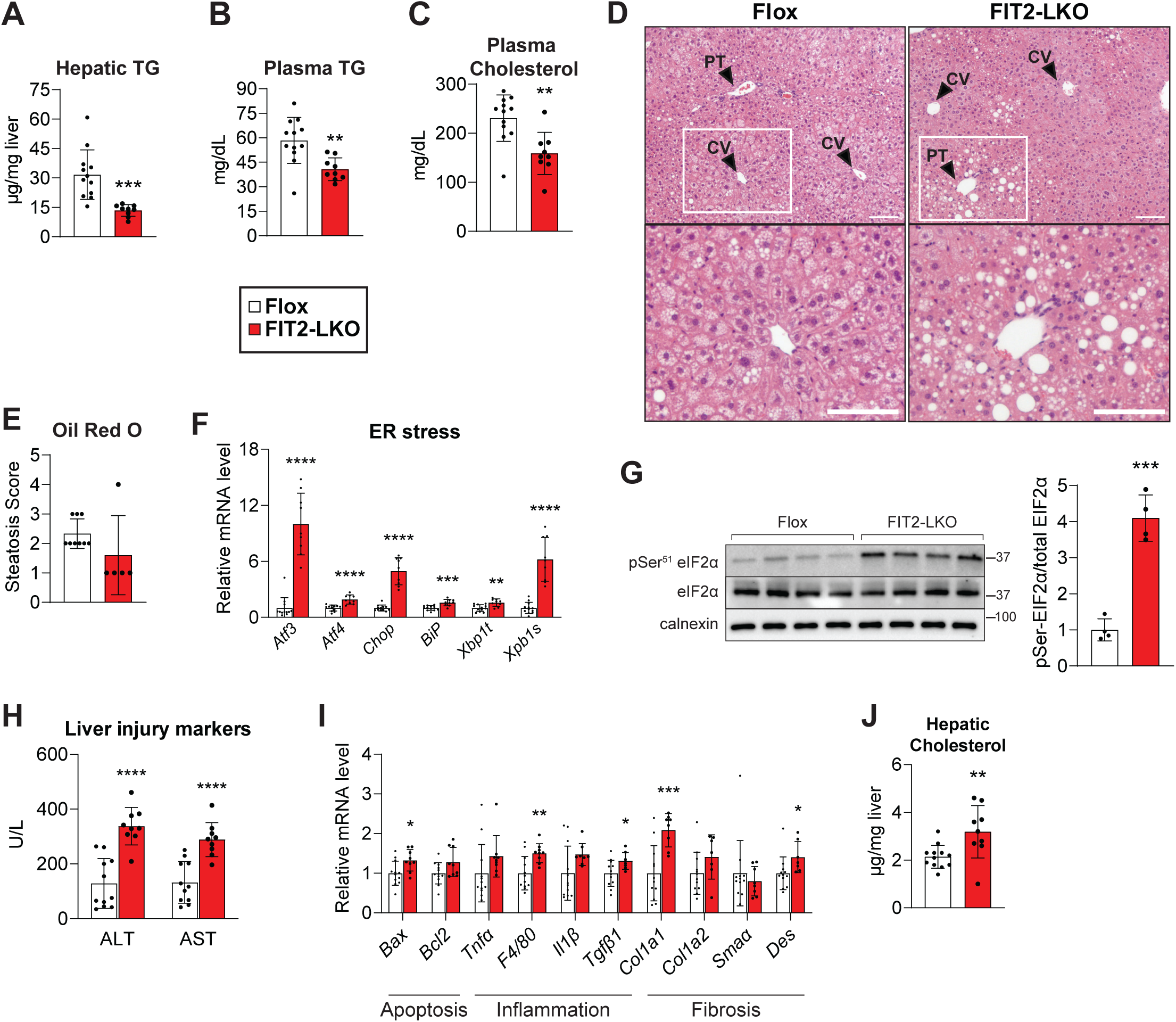
High-fat-diet feeding exacerbates ER stress and liver injury in FIT2 liver-specific KO mice. Levels of hepatic TG (A), plasma TG (B), and plasma cholesterol (C) are decreased in FIT2-LKO mice after weeks high-fat diet (HFD) feeding. (D) Representative images of H&E staining of livers from Flox and 2-LKO mice after HFD feeding; CV=central vein, PT=portal triad. Scale bar=50 µm. (E) Steatosis scoring Oil Red O staining of Flox and FIT2-LKO mice given HFD challenge (n=5-9/genotype). (F) RT-qPCR ies show that under HFD, FIT2-LKO mice have increased expression of ER stress genes. (G) This is supported by increased phosphorylation of the UPR protein eIF2α, as shown with western blotting 4/genotype). (H) After HFD challenge, FIT2-LKO exhibit exacerbated liver injury, as shown by measurement of plasma alanine transaminase and aspartate transaminase (ALT and AST). (I) This phenotype ents with relatively minor changes in the expression of gene markers for apoptosis, inflammation, or osis. (J) FIT2-LKO mice had increased levels of hepatic cholesterol. Data represents mean ± SD. 9-12/genotype, unless otherwise noted above for specific experiments. Statistical significance was luated with unpaired Student’s 2-tailed t-test for (A-C, G, J). For (E, F, I), statistical significance was luated with unpaired Student’s 2-tailed t-test for parametric data and a Mann-Whitney U test for parametric data. For (H), 2-way ANOVA with Šidák correction was used. *p<0.05, **p<0.01, ***p<0.001, ****p<0.0001.

As with chow feeding, FIT2-LKO mice showed evidence of hepatic ER stress and injury. Levels of both mRNA transcripts and phosphorylated protein markers of the unfolded protein response were increased in the FIT2-LKO mice to an even greater degree than under standard chow feeding (Figure 4F-G). Moreover, plasma ALT and AST markers of liver injury were further increased in the FIT2-LKO mice after HFD challenge (Figure 4H). Though apoptotic, inflammation, and fibrosis often accompany such severe ER stress and liver damage, these markers were not substantially altered between genotypes (Figure 4I).

## Discussion

Previous data showed that the ER-resident FIT2 has acyl-CoA diphosphatase activity in vitro and is important for maintaining ER homeostasis in human and yeast cells (14). We now show that hepatic deficiency of FIT2 results in increased acyl-CoA levels in vivo, which are linked to increased ER stress and signs of liver injury, as manifested by elevated circulating transaminase levels. These findings for hepatic FIT2 deficiency were exacerbated with HFD feeding. Although it is uncertain whether humans with FIT2 deficiency exhibit similar hepatocyte defects (10,11), our studies highlight the crucial importance of FIT2 in lipid and ER homeostasis in vivo.

In contrast to what was found in FIT2-deficient cultured cells (14), we found no gross morphological changes of the ER or ER whorls in hepatocytes of FIT2-LKO mice. A possible explanation for the absence of ER morphology changes could be that increased autophagic flux cleared such structures, particularly since autophagy ameliorates liver damage in certain contexts (21). Consistent with this notion, FIT2 interacts genetically with autophagic pathways; FIT2 deletion sensitizes Renca cancer cells to cell death from IFNγ, and inactivation of autophagy reverses this phenotype (12).

The reduction in TG storage under HFD feeding conditions is similar to what has been reported for FIT2 deficiency in cells that have been cultured with excess fatty acids (1,14). However, unexpectedly, chow-fed FIT2-LKO mice accumulated neutral lipids and TGs in hepatocytes. The modest level of steatosis in chow-fed FIT2-LKO mice may be at least partially explained by reductions in TG secretion and fatty acid oxidation capacity. With respect to TG secretion, our results are consistent with the hypothesis that FIT2 deficiency impairs loading of nascent lipoproteins with TGs. Since FIT2 is hypothesized to act on the lumenal leaflet of the ER, FIT2 deficiency may lead to acyl-CoA accumulation at this leaflet and interfere with lipidation of the nascent apoB particles in the ER lumen. Similarly, changes in ER phospholipids can have marked effects on TG secretion (17,22,23).

The reduction in fatty oxidation capacity in lysates of the FIT2-LKO livers was substantial and may also contribute to the TG accumulation in chow-fed mice. We found no differences in mitochondrial content, gene expression, or protein levels, suggesting that FIT2 deficiency adversely affects fatty acid oxidation through an as-yet unknown mechanism. Of note, altered mitochondrial biology is consistent with the human FIT2 deficiency phenotype; patients with FIT2 mutations present with deafness-dystonia symptoms similar to those afflicted with Mohr-Tranebjaerg syndrome, which is caused by defects in mitochondrial function (19,20).

Our findings highlight that the phenotypes of ER stress and lipid accumulation with FIT2 deficiency can be dissociated. We found ER stress to be a consistent observation in all our studies of FIT2 deficiency in cells and mice, on either diet. In contrast, the lipid accumulation phenotype in mice appears to be contextual and depends on the dietary status. This supports the hypothesis that the lipid storage phenotypes are a secondary consequence and not a primary role for FIT2 in LD formation (14). In support of this, loss of FIT2 in pancreatic β cells is accompanied by ER stress, and FIT2 deficiency resulted in several-fold increased levels of tissue ceramides (9). We also found ceramide levels were increased in FIT2-LKO livers, but to a lesser extent (∼30%) (Figure 1K).

The mechanism for how FIT2 deficiency results in ER stress is unclear. Most proximally, the defects associated with FIT2 deficiency are likely due either to accumulation of its substrates (e.g., unsaturated acyl-CoAs) or to deficiency of its products (i.e., 3′,5′-ADP and acyl 4′-phosphopantetheine). Although this remains uncertain at present, FIT2 activity is clearly important for the health of cells. Interestingly, human FIT2 protects Renca cancer cells from IFNγ effects (12). Additionally, high intra-tumoral levels of *FITM2* expression correlate with decreased survival in human patients with hepatocellular carcinoma (3). Thus, FIT2 inhibitors may be useful to sensitize specific cancer cells to targeted chemotherapies. Continued studies to elucidate the consequences of FIT2 deficiency will be essential for unraveling why FIT2 activity is so crucial to cell health and, hopefully, for finding therapies for humans suffering from the consequences of FIT2 deficiency.

## Acknowledgments

We thank members of the Farese & Walther laboratory and D. Silver for helpful discussions and D. Silver for sharing reagents; M. Becuwe for contribution to early mouse line generation; the Rodent Histopathology Core at Dana Farber/Harvard Cancer Center in Boston, Massachusetts, supported, in part, by an NCI Cancer Center Support Grant #NIH5P30 CA06516, which provided tissue sectioning and staining services; and the Electron Microscopy Facility at Harvard Medical School for electron microscopy sample preparation and microscopy training and access. This work was supported by R01GM141050 (to R.V.F). L.M.B. was supported by the National Institute of Health Service Award T32 DK00747. L.M.B. and A.I. were supported by American Heart Association Postdoctoral Fellowships. R.L.W.’s efforts were supported by the Institute for Advancing Health through Agriculture and Texas AgriLife Research project #8738. T.C.W. is an investigator of the Howard Hughes Medical Institute.

## Author Contributions

L.M.B., R.V.F., and T.C.W., conceived the project. L.M.B., R.V.F., and T.C.W. designed the experiments. L.M.B. and A.I. performed and analyzed most of the experiments. Z.W.L. performed the lipidomics and proteomics experiments. R.L.W. performed the dynamic light scattering experiment. O.I. conducted acyl-CoA measurements. R.B. performed histological analyses and scoring. L.M.B., A.I., R.V.F., and T.C.W. wrote the manuscript. All authors discussed the results and contributed to the manuscript.

## TABLES

**Table S1**. Mass spectrometry measurement of FIT1 and FIT2 in liver and skeletal muscle.

**Table S2. Hepatic acyl-CoA measurements**. Absolute and relative amounts of CoA and acyl-CoA species in Flox and FIT2-LKO livers, as measured by mass spectrometry. n=6/genotype. *p<0.05.

**Table S3**. qPCR primer sequences.

## MATERIALS AND METHODS

### Animals

Fitm2 ^*flox/flox*^ mice (028832, Jackson Laboratory, Miranda et al., 2014) were crossed with mice expressing Cre recombinase under control of the albumin promoter (B6.Cg-Tg(Alb-Cre)21MGN/J). Mice were housed in the Harvard School of Public Health animal care facility and maintained on a 12-h light-dark cycle. Mice had free access to water and food unless specified otherwise. Mice were weaned to a standard rodent chow diet (PicoLab Rodent Diet 20 #5053, St. Louis, MO). For high-fat diet studies, animals were weaned to a chow diet and then fed TD.88137 (42% calories from fat) for 11 weeks starting at 7 weeks of age. For food intake measurements, animals were individually caged 1 week prior to data collections. All mice were male and fasted 2 hours prior to sacrifice unless specified otherwise. Mice were euthanized with isoflurane, blood was collected via cardiac puncture, and tissues were collected. All *in-vivo* mouse experiments were conducted in accordance with protocol approved by the Institutional Animal Care and Use Committee at the Harvard University.

### Quantitative PCR

Tissues were homogenized in Qiazol using a Bead Mill Homogenizer (VWR), RNA was isolated using RNeasy kit (Qiagen), and cDNA was synthesized using an iSCRIPT cDNA synthesis kit (Bio-Rad). qPCR was performed using Power SYBR Green PCR Master Mix kit (Applied Biosystems). Primers sequences are listed in Table S3.

### Immunoblotting

Livers were homogenized in RIPA buffer (Cell Signaling Technology), supplemented with complete mini EDTA-free protease inhibitor (Sigma Aldrich) and PhosSTOP phosphatase inhibitor (Sigma-Aldrich) using a Bead Mill Homogenizer (VWR). Protein concentrations were measured using a DC Protein Assay (Bio-Rad). Proteins were incubated at 60°C for 15 min in 4x Laemmli sample buffer (Bio-Rad). 20–40µg liver lysate protein was separated by SDS-PAGE and transferred to a PVDF membrane. The membrane was blocked with TBS-T containing 5% nonfat dry milk for 1 h at room temperature and then incubated overnight at 4°C in primary antibody. Membranes were washed in TBS-T, incubated in secondary antibody, washed with TBS-T and visualized using SuperSignal Chemiluminescent Substrate (Thermo Scientific). Band intensity was measured using ImageJ software. Antibodies used include: total eIF2a (Cell Signaling Technology #9722, 1:1000, in 5% BSA), phosphor-Ser51-eIF2a (Cell Signaling Technology #9721, 1:1000),vinculin (Cell Signaling Technology #4650, 1:1000), ApoB (Abcam ab31992, 1:1000), and FIT2 (a generous gift from David Silver’s laboratory (1), 1:1000)

### Acyl-CoA measurements

Cellular and liver acyl-CoA esters were analyzed using a method based on a report by Magnes et al. (24) that relies on the extraction procedure described by Deutsch et al. (25). The CoAs were further purified by solid phase extraction as described by Minkler et al. (26). The acyl CoAs were analyzed by flow injection analysis using positive electrospray ionization on Xevo TQ-S, triple quadrupole mass spectrometer (Waters, Milford, MA) employing methanol/water (80/20, v/v) containing 30 mM ammonium hydroxide as the mobile phase. Spectra were acquired in the multichannel acquisition mode monitoring the neutral loss of 507 amu (phosphoadenosine diphosphate) and scanning from m/z 750-1060. Heptadecanoyl CoA was employed as an internal standard. The endogenous CoAs were quantified using calibrators prepared by spiking cell or liver homogenates with authentic CoAs (Sigma, St. Louis, MO) having saturated acyl chain lengths C_0_-C_18._. Corrections for the heavy isotope effects, mainly ^13^C, to the adjacent m+2 spectral peaks in a particular chain-length cluster were made empirically by referring to the observed spectra for the analytical standards.

### Proteomics

Liver (∼20 mg) was homogenized in 800 μl of PBS (supplemented with complete mini EDTA-free protease inhibitor (Sigma Aldrich) and 5 mM EDTA) using a Bead Mill Homogenizer (VWR). Extraction of proteins were performed as described (27). Mass spectrometry data were analyzed by MaxQuant software version 1.5.2.8 (28) using the following setting: oxidized methionine residues and protein N-terminal acetylation as variable modification, cysteine carbamidomethylation as fixed modification, first search peptide tolerance 20 ppm, and main search peptide tolerance 4.5 ppm. Protease specificity was set to trypsin with up to two missed cleavages allowed. Only peptides longer than six amino acids were analyzed, and the minimal ratio count to quantify a protein is 2. The false discovery rate was set to 5% for peptide and protein identifications. Database searches were performed using the Andromeda search engine integrated into the MaxQuant environment (29) against the UniProt-mouse database containing 54,185 entries (December 2018). “Matching between runs” algorithm with a time window of 0.7 min was utilized to transfer identifications between samples processed using the same nanospray conditions. Protein tables were filtered to eliminate identifications from the reverse database and common contaminants. Fold changes of proteins were calculated comparing mean area of log2 intensities between replicates of different genotypes. Statistical significance was calculated using a Student’s t-test followed by Benjamini-Hochberg FDR correction of 5% for multiple hypothesis testing. The mass spectrometry proteomics data have been deposited to the ProteomeXchange Consortium via the PRIDE partner repository (30) with the dataset identifier PXD033884.

### Lipidomics

Liver (∼100mg) was homogenized in 1 mL of PBS using a Bead Mill Homogenizer (VWR). Lipids were extracted, according to the Folch method (31). Lysis volume was normalized to starting tissue material. Organic fraction containing extracted lipids was subjected to LC-MS/MS analysis, as described in (27). Mass spectrometry data analysis was performed using LipidSearch version 4.1 SP (Thermo Fisher Scientific). The results were exported to R-Studio where quality control was performed using pairwise correlations between replicates, a principal component analysis comparing sample groups, as well as retention time plot analysis to verify elution clustering within lipid classes. All identified lipids were included for subsequent analyses if they fulfilled the following LipidSearch-based criteria: 1) reject equal to zero, 2) main grade A OR main grade B AND APvalue<0.01 for at least three replicates, and 3) no missing values across all samples. S Statistical significance was calculated using a Student’s t-test followed by a Holm-Sidak test to correct for multiple comparisons.

### Histology

Livers were collected and fixed in formalin overnight at 4°C. Livers were sectioned and stained by the Rodent Histopathology Core at Harvard Medical School. Frozen sections were used for Oil Red O staining, which were unbiasedly scored for steatosis by a histopathologist (27). Paraffin-embedded tissue was used for H&E staining. Sections were imaged on a ZEISS light microscope.

### Electron microscopy

Mice were anesthetized with isoflurane and then perfused with 10 mL of PBS followed by 10 mL of 2.5% glutaraldehyde, 2.5% paraformaldehyde in 0.1 M sodium cacodylate buffer (pH 7.4). 1–2-mm liver pieces were fixed in the fixative overnight, washed several times in 0.1M cacodylate buffer, osmicated in 1% osmium tetroxide/1.5% potassium ferrocyanide (final solution) for 3 h, followed by several washes of dH_2_O. 1% uranyl acetate in maleate buffer was added for 1 h and then washed several times with maleate buffer (pH 5.2). This was followed by a graded cold ethanol series up to 100%, which is changed 3x over 1 h, followed by propylene oxide, changed 3x over 1 h. The sample was then placed in ½ and ½ propylene oxide with plastic mixture including catalyst overnight. The following day, samples were polymerized in Taab 812 Resin (Marivac Ltd., Nova Scotia, Canada) at 60° for 24-48 h. 80-nm sections were cut with a Leica ultracut microtome, picked up on 100 mesh formvar/carbon coated copper grids, stained with 0.2% Lead Citrate, and viewed and imaged with a JEOL 1200X electron microscope equipped with an MP 2k CCD camera. For ER dilation quantification, three images (representative of ER dilation for that animal) were selected per mouse, and the distance across the ER lumen (bilayer center-to-bilayer center) was measured using ImageJ. At least 25 lumen measurements were calculated per image and averaged to provide the representative ER dilation for that mouse. For total ER quantification, three images (representative of ER content for that a) were selected per mouse. Using ImageJ, the total cell area was traced and calculated (nucleus excluded due to variability in nuclear size), and the ER was manually traced. ER content was calculated as nm ER length divided by µm^2^ available cell area. Four flox and seven FIT2-LKO animals were assessed using this method, and the data depict the average and standard deviation of these biological samples.

### TG and apoB secretion measurements

Mice were fasted for 4 h and injected intravenously with 1000 mg/kg body weight with the lipoprotein lipase inhibitor, Polaxomer-407. Tail vein blood was collected at t=0, 1, 2, and 4 h. Plasma was supplemented with complete mini EDTA-free protease inhibitor (Sigma Aldrich) and snap frozen. Plasma TG was measured using Infinity TG kit (Thermo Scientific). ApoB-100 protein levels were measured by immunoblotting, as described above. At 1 h, the sample was diluted 1:5. At 2 and 4 h, they were diluted 1:10. 1.5 µL of plasma (or 1.5 µL of diluted plasma) were heated at 95° for 5 min in 2x denaturing sample buffer. Plasma samples from each time point were run on the same gel. Equal loading was confirmed by visualization of albumin with Ponceau staining. To compare between gels, a sample from each time point was run on the t=0 gel (Figure S4C).

### VLDL particle size measurements

Mice were fasted 4 h prior to sacrifice. 2 × 10 µl removed for density profiling by isopycnic ultracentrifugation (32). For each plasma sample, the d < 1.063 g/ml fraction was prepared by ultracentrifugation in a Beckman Coulter Optima MAX-XP benchtop ultracentrifuge in an MLA-55 rotor (18h X 172,301 x G at 14°C). This fraction contains virtually all of the apo-B100 in the plasma (33). Lipoprotein particle diameters were determined by dynamic light scattering analysis with a Microtrac Series 150 Ultrafine particle analyzer fitted with a flexible conduit-sheathed probe tip (UPA-150; Microtrac, Clearwater, Florida, USA) (34,35). Raw particle-size distributions from number distributions were converted to population percentiles, which were used to calculate the median particle diameter for each decile of lipoprotein size distribution.

### Ex vivo fatty acid oxidation assay

Mice were fasted for 4 h and euthanized with isoflurane. Liver was collected and processed via Dounce homogenization in 2 mL of sucrose-Tris-EDTA buffer as detailed (36). Liver homogenates were centrifuged at 450xg for 10 min at 4ºC. Supernatants were collected and incubated for 1 h in the presence of fatty acid oxidation substrate (300 μM palmitic acid with 0.4 μCi 1-14C-palmitic acid). 1 mM rotenone, an inhibitor of oxidative phosphorylation, was used as a control. Radioactivity of trapped CO_2_ and acid-soluble metabolites were measured using a liquid scintillation counter. Fatty acid oxidation rates were calculated as [(counts per minute-blank)/reaction mixture specific activity]/gram tissue.

### Liver biochemical assays

Liver (∼50 mg) was homogenized in 500 μL of lysis buffer (250 mM sucrose, 50 mM Tris HCl, pH 7.4). Lipids were extracted using a modified Bligh and Dyer method and solubilized in 0.1% Triton-X-100 by sonication (three rounds of 2 s at 30 mA). TG and cholesterol were quantified using Infinity Triglyceride and Cholesterol Reagents (Thermo Scientific). Glycogen was measured from ∼20 mg of liver using EnzyChrom™ Glycogen Assay Kit (BioAssay Systems), according to the manufacturer’s instructions.

### Plasma analyses

Plasma TG and total plasma cholesterol were measured from 2 and 10 µL of plasma, respectively, using Infinity Triglyceride and Cholesterol Reagents (Thermo Scientific). For HDL cholesterol measurements, non-HDL was precipitated by incubating 20 µL of plasma with precipitation buffer containing 0.44 mM phosphotungstic acid and 20 mM MgCl_2_ for 10 min at room temperature, followed by centrifugation. Cholesterol was measured from the resulting supernatant. Plasma ALT, AST, bilirubin, ALP, albumin and total protein were measured with Piccolo Liver Panel Plus discs used with a Piccolo Xpress chemistry analyzer (Abaxis).

### Statistical analyses

Results are expressed as mean ± standard deviation. Statistical significance was evaluated with unpaired Student’s 2-tailed t-test (if data passed a Shapiro-Wilk test for normality) or a Mann-Whitney U test (for nonparametric data, which did not pass test for normality). For experiments with multiple readouts, statistical significance was evaluated with two-way analysis of variance (ANOVA) with post-hoc Šidák test, or repeated-measures ANOVA for time-course experiments. Analyses were performed using GraphPad Prism 7. *p<0.05, **p<0.01, ***p<0.001, ****p<0.0001.

### Study Approval

All in vivo mouse experiments were conducted in accordance with protocol approved by the Institutional Animal Care and Use Committee at the Harvard University.

