## Supplemental Material for "*Fitm2* is required for ER homeostasis and normal function of murine liver"

**Table S1.** Mass spectrometry measurement of FIT1 and FIT2 in liver and skeletal muscle.

| ID | Flox 1 | Flox 2 | Flox 3 | Flox 4 | FIT2-LKO 1 | FIT2-LKO 2 | FIT2-LKO 3 | FIT2-LKO 4 | FIT2-LKO 5 | Flox 1 |
|---|---|---|---|---|---|---|---|---|---|---|
| Tissue | Liver | Liver | Liver | Liver | Liver | Liver | Liver | Liver | Liver | Skeletal Muscle |
| FITM1 | n.d. | n.d. | n.d. | n.d. | n.d. | n.d. | n.d. | n.d. | n.d. | 26.0766 |
| FITM2 | 26.4376 | 23.3978 | 21.1139 | 21.6667 | n.d. | n.d. | n.d. | n.d. | n.d. | n.d. |
| ACTB | 33.1945 | 34.6911 | 33.1693 | 32.9812 | 33.3333 | 32.6851 | 32.7691 | 33.077 | 33.0558 | 29.4254 |
| FASN | 33.8659 | 34.3608 | 33.6764 | 33.347 | 34.1397 | 33.5568 | 33.5418 | 33.7423 | 34.2724 | 24.6743 |
| MYH2 | n.d. | n.d. | n.d. | n.d. | n.d. | n.d. | n.d. | n.d. | n.d. | 33.2602 |
| MYH3 | n.d. | n.d. | n.d. | n.d. | n.d. | n.d. | n.d. | 20.3023 | n.d. | 32.9784 |

**Table S2. Hepatic acyl-CoA measurements.** Absolute and relative amounts of CoA and acyl-CoA species in Flox and FIT2-LKO livers, as measured by mass spectrometry. n=6/genotype. *q<0.05.

| | nmol/g liver | | | | relative abundance | | | | q-value |
|---|---|---|---|---|---|---|---|---|---|
| | AVE | | SD | | AVE | | SD | | |
| | Flox | LKO | Flox | LKO | Flox | LKO | Flox | LKO | |
| **free CoA** | 65.3 | 57.8 | 6.67 | 6.23 | 1.00 | 0.89 | 0.10 | 0.10 | 0.123 |
| **acetyl** | 79.5 | 71.2 | 10.7 | 6.21 | 1.00 | 0.89 | 0.14 | 0.08 | 0.191 |
| **propionyl** | 3.42 | 2.75 | 0.95 | 0.64 | 1.00 | 0.80 | 0.28 | 0.19 | 0.211 |
| **crotonyl** | 1.00 | 1.01 | 0.15 | 0.08 | 1.00 | 1.01 | 0.15 | 0.08 | 0.788 |
| **butyryl** | 5.64 | 6.70 | 1.01 | 1.12 | 1.00 | 1.19 | 0.18 | 0.20 | 0.174 |
| **b-hydroxy propionyl** | 0.47 | 0.51 | 0.06 | 0.08 | 1.00 | 1.09 | 0.13 | 0.17 | 0.788 |
| **3-me-crotonyl** | 0.53 | 0.52 | 0.17 | 0.14 | 1.00 | 0.98 | 0.31 | 0.27 | 0.788 |
| **isovaleryl** | 1.44 | 1.45 | 0.50 | 0.41 | 1.00 | 1.01 | 0.35 | 0.29 | 0.783 |
| **malonyl/hydroxybutyryl** | 7.62 | 9.01 | 1.92 | 1.23 | 1.00 | 1.18 | 0.25 | 0.16 | 0.204 |
| **succinyl** | 0.41 | 0.47 | 0.08 | 0.11 | 1.00 | 1.16 | 0.20 | 0.26 | 0.428 |
| **itaconyl** | 0.31 | 0.46 | 0.14 | 0.16 | 1.00 | 1.48 | 0.45 | 0.53 | 0.196 |
| **glutaryl** | 0.66 | 0.97 | 0.16 | 0.13 | 1.00 | 1.46 | 0.25 | 0.19 | 0.015* |
| **hexenoyl** | 0.48 | 0.60 | 0.10 | 0.08 | 1.00 | 1.25 | 0.21 | 0.17 | 0.259 |
| **hexanoyl** | 1.97 | 3.12 | 0.52 | 0.78 | 1.00 | 1.59 | 0.26 | 0.40 | 0.027* |
| **8:00** | 0.84 | 1.44 | 0.21 | 0.48 | 1.00 | 1.71 | 0.25 | 0.57 | 0.034* |
| **10:0** | 0.81 | 0.87 | 0.19 | 0.17 | 1.00 | 1.07 | 0.23 | 0.21 | 0.608 |
| **12:1** | 0.37 | 0.31 | 0.13 | 0.04 | 1.00 | 0.84 | 0.34 | 0.11 | 0.555 |
| **12:0** | 0.49 | 0.40 | 0.15 | 0.07 | 1.00 | 0.81 | 0.30 | 0.15 | 0.264 |
| **14:2** | 1.15 | 0.82 | 0.55 | 0.20 | 1.00 | 0.72 | 0.48 | 0.18 | 0.264 |
| **14:0** | 1.21 | 1.24 | 0.35 | 0.14 | 1.00 | 1.03 | 0.29 | 0.11 | 0.783 |
| **14:1** | 1.02 | 1.13 | 0.50 | 0.29 | 1.00 | 1.11 | 0.49 | 0.29 | 0.608 |
| **16:0** | 4.84 | 7.00 | 0.93 | 1.20 | 1.00 | 1.45 | 0.19 | 0.25 | 0.017* |
| **16:1** | 4.33 | 6.08 | 0.73 | 1.81 | 1.00 | 1.41 | 0.17 | 0.42 | 0.098 |
| **18:0** | 2.99 | 5.01 | 0.60 | 0.49 | 1.00 | 1.68 | 0.20 | 0.17 | 0.001* |
| **18:1** | 8.70 | 19.86 | 1.23 | 5.05 | 1.00 | 2.28 | 0.14 | 0.58 | 0.002* |
| **18:2** | 8.92 | 9.94 | 1.05 | 1.24 | 1.00 | 1.12 | 0.12 | 0.14 | 0.204 |
| **18:3** | 1.52 | 1.25 | 0.14 | 0.25 | 1.00 | 0.82 | 0.09 | 0.17 | 0.098 |
| **19:0** | 0.95 | 1.60 | 0.26 | 0.25 | 1.00 | 1.68 | 0.28 | 0.26 | 0.005* |
| **19:1** | 0.70 | 0.87 | 0.19 | 0.11 | 1.00 | 1.24 | 0.27 | 0.15 | 0.123 |
| **20:0** | 1.27 | 1.73 | 0.32 | 0.11 | 1.00 | 1.36 | 0.25 | 0.09 | 0.017* |
| **20:1** | 1.84 | 5.85 | 0.47 | 1.48 | 1.00 | 3.18 | 0.26 | 0.80 | 0.001* |
| **20:2** | 1.49 | 3.16 | 0.51 | 0.40 | 1.00 | 2.12 | 0.34 | 0.27 | 0.001* |
| **20:3** | 0.95 | 1.50 | 0.14 | 0.15 | 1.00 | 1.57 | 0.15 | 0.16 | 0.001* |
| **20:4** | 2.74 | 2.56 | 0.29 | 0.35 | 1.00 | 0.93 | 0.10 | 0.13 | 0.377 |
| **20:5** | 1.21 | 1.83 | 0.11 | 0.22 | 1.00 | 1.51 | 0.09 | 0.18 | 0.001* |
| **21:0** | 0.36 | 0.66 | 0.08 | 0.10 | 1.00 | 1.82 | 0.23 | 0.26 | 0.001* |
| **21:1** | 0.26 | 0.37 | 0.05 | 0.06 | 1.00 | 1.40 | 0.18 | 0.24 | 0.017* |
| **22:1** | 3.21 | 3.45 | 0.91 | 0.80 | 1.00 | 1.07 | 0.28 | 0.25 | 0.608 |
| **22:2** | 0.80 | 1.21 | 0.16 | 0.24 | 1.00 | 1.51 | 0.20 | 0.30 | 0.017* |
| **22:5** | 0.45 | 0.60 | 0.05 | 0.05 | 1.00 | 1.32 | 0.11 | 0.12 | 0.002* |
| **22:6** | 1.61 | 1.56 | 0.17 | 0.13 | 1.00 | 0.97 | 0.11 | 0.08 | 0.759 |

**Table S3.** qPCR primer sequences.

| Gene name | F/R | Sequence |
|---|---|---|
| 18S | F | CGCTTCCTTACCTGGTTGAT |
| 18S | R | GAGCGACCAAAGGAACCATA |
| Acly | F | AAAGCTTGGCCTCGTCGG |
| Acly | R | GGGACGAAGGGTTCAATGAGA |
| Aox | F | CTCATCTTCGAGGCTTGGAAACCAC |
| Aox | R | ATTTCACGGATAGGGACAAGAAAGGC |
| Arbp | F | TCACTGTGCCAGCTCAGAAC |
| Arbp | R | AATTTCAATGGTGCCTCTGG |
| Atf3 | F | GAGGATTTTGCTAACCTGACACC |
| Atf3 | R | TTGACGGTAACTGACTCCAGC |
| Atp5a1 | F | TCCATGCCTCTAACACTCGAC |
| Atp5a1 | R | GACGTGTCAGCTCCCAGAA |
| Bax | F | GGAGATGAACTGGACAGCAATA |
| Bax | R | GAAGTTGCCATCAGCAAACAT |
| Bcl2 | F | GAGCAGGTGCCTACAAGAAA |
| Bcl2 | R | CTTTGTCCTCTGACTGGGTATG |
| Bip | F | ACTTGGGGACCACCTATTCCT |
| Bip | R | ATCGCCAATCAGACGCTCC |
| Chop | F | CCACCACACCTGAAAGCAGAA |
| Chop | R | AGGTGAAAGGCAGGGACTCA |
| Chrebpa | F | CGACACTCACCCACCTCTTC |
| Chrebpa | R | TTGTTCAGCCGGATCTTGTC |
| Chrebpb | F | TCTGCAGATCGCGTGGAG |
| Chrebpb | R | CTTGTCCCGGCATAGCAAC |
| Co1 | F | CCCAATCTCTACCAGCATC |
| Co1 | R | GGCTCATAGTATAGCTGGAG |
| Cox4i1 | F | TCACTGCGCTCGTTCTGAT |
| Cox4i1 | R | CGATCGAAAGTATGAGGGATG |
| Cpt1a | F | GAA CCC CAA CAT CCC CAA AC |
| Cpt1a | R | TCC TGG CAT TGT CCT GGA AT |
| Cre | F | CGGTCTGGCAGTAAAAACTAT |
| Cre | R | CAGGGTGTTATAAGCAATCCC |
| Cre | F | CCCTGTTTCACTATCCAGGT |

| | | |
|---|---|---|
| Cre | R | GGGTAACTAAACTGGTCGAG |
| Cyclo | F | TGGAGAGCACCAAGACAGACA |
| Cyclo | R | TGCCGGAGTCGACAATGAT |
| Dgat1 | F | GGAATATCCCCGTGCACAA |
| Dgat1 | R | CATTTGCTGCTGCCATGTC |
| Dgat2 | F | CCGCAAAGGCTTTGTGAA |
| Dgat2 | R | GGAATAAGTGGGAACCAGATCAG |
| Edem | F | AAGCCCTCTGGAACCTTGCG |
| Edem | R | AACCCAATGGCCTGTCTGGT |
| F4/80 | F | TGACTCACCTTGTGGTCCTAA |
| F4/80 | R | CTTCCCAGAATCCAGTCTTTCC |
| Fitm2 | F | CGAGAGCTACCTCAGCAACAAG |
| Fitm2 | R | CAATGAAAGGCAGGAGGAGACA |
| H19 | F | GTACCCACCTGTCGTCC |
| H19 | R | GTCCACGAGACCAATGACTG |
| Hmgcr | F | CTTGTGGAATGCCTTGTGATTG |
| Hmgcr | R | AGCCGAAGCAGCACATGAT |
| Hmgcs | F | GCCGTGAACTGGGTCGAA |
| Hmgcs | R | GCATATATAGCAATGTCTCCTGCAA |

#### Figure S1

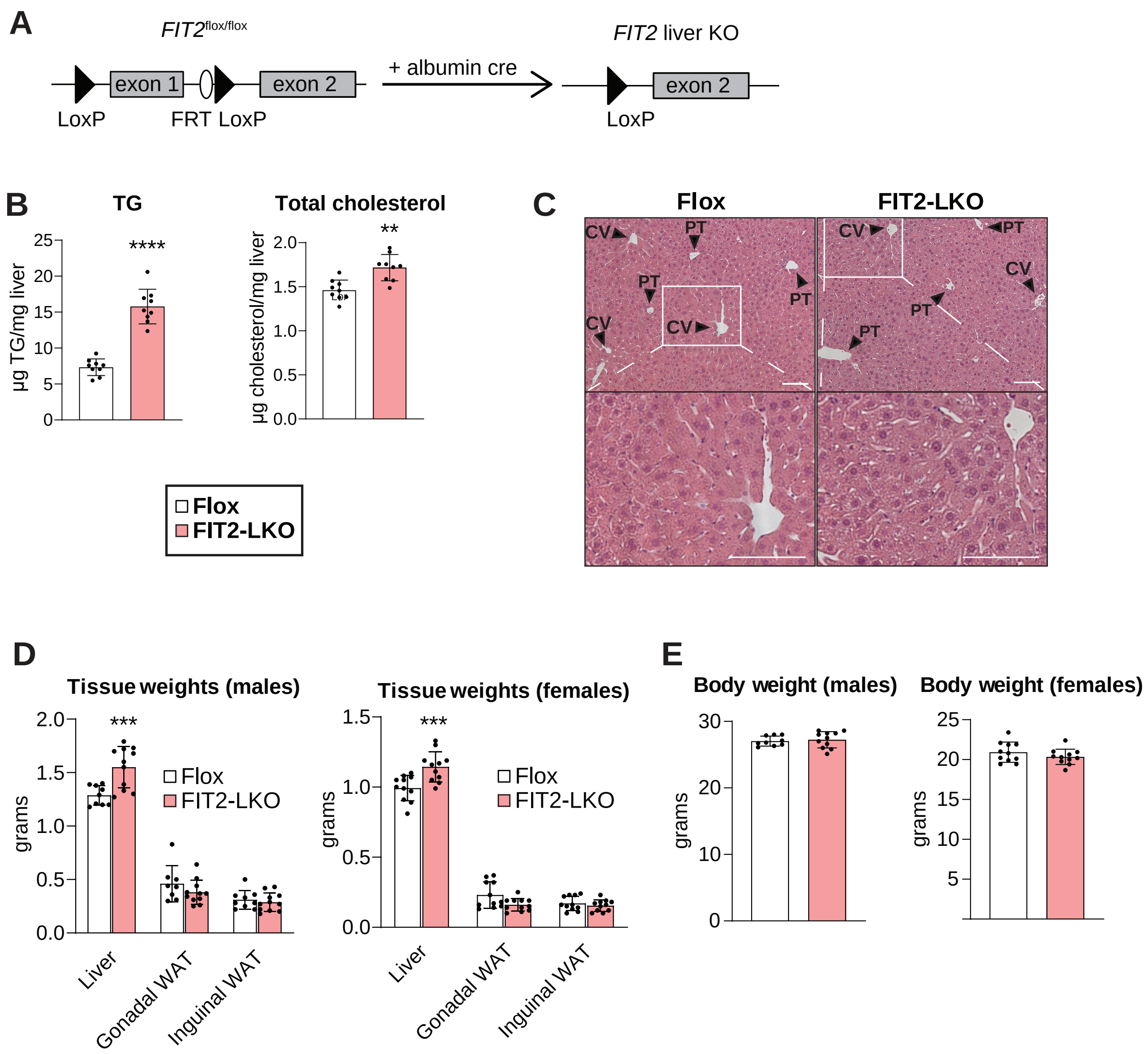


**Figure S1. Corresponding to Figure 1.** (A) Genetic construct used for generation of murine model of FIT2 deficiency in hepatocytes. (B) Liver-specific FIT2 deficiency elevates liver TG and total cholesterol in female mice (n=9/genotype). (C) Representative images of H&E staining of livers from male Flox and FIT2-LKO mice. Scale bar=50 µm. CV=central vein; PT=portal triad. (D) FIT2-LKO mice have heavier livers than flox littermates. (E) FIT2-LKO mice do not exhibit altered body weight, compared to flox control littermates with chow feeding (n=9-11/genotype). Data represents mean ± SD. Statistical significance was evaluated with unpaired Student's 2-tailed t-test (B, E) and 2-way ANOVA with Šidák correction in (D). *p<0.05, **p<0.01, ***p<0.001, ****p<0.0001.

#### Figure S2

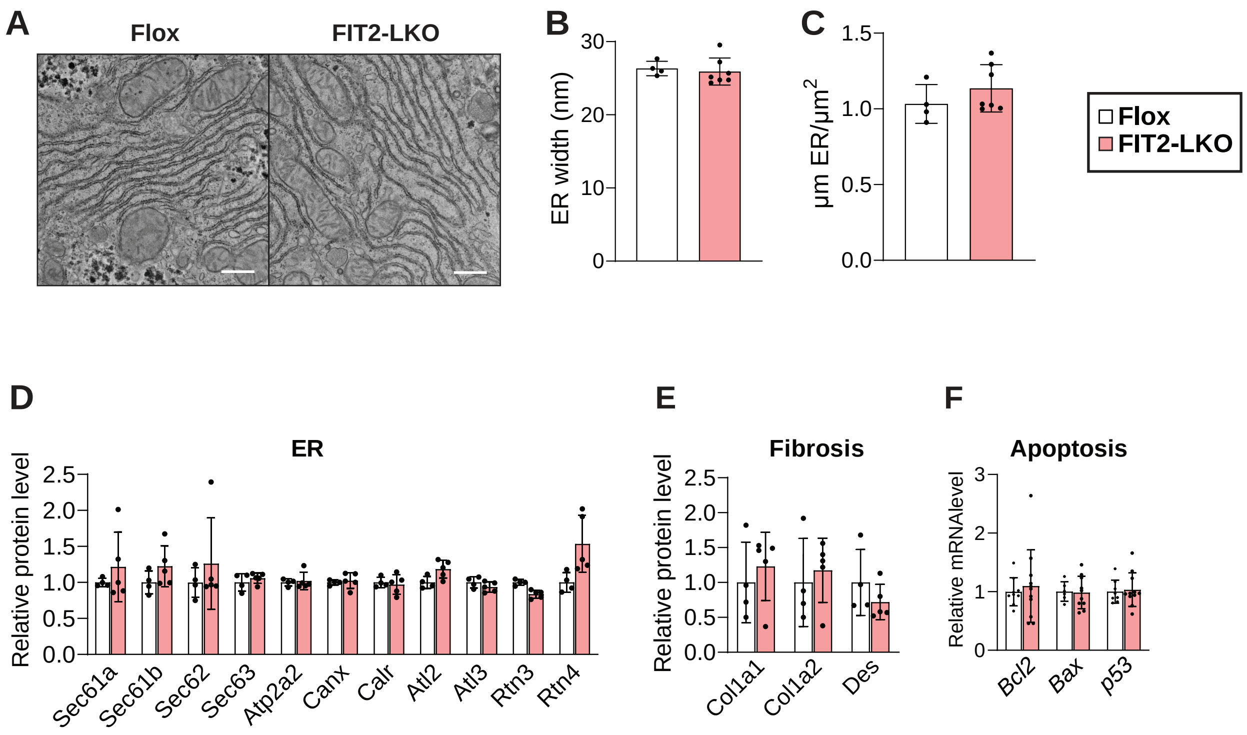


**Figure S2. Corresponding to Figure 2.** Representative electron microscopy images (A) and quantification of ER dilation (B) indicate that loss of FIT2 does not alter ER ultrastructure in hepatocytes (n=5–8). Flox and FIT2-LKO hepatocytes contain equal amounts of ER, as quantified from electron microscopy images (n=5–8/genotype) (C). Protein abundance of ER proteins (D) and fibrosis markers (E), as measured by mass spectrometry (n=4–5/genotype). (F) FIT2-LKO mice do not exhibit evidence for apoptosis; transcript levels of anti-apoptotic (Bcl2) and pro-apoptotic (Bax, p53) gene markers are unaltered (n=9–11/genotype). Data represents mean ± SD. Statistical significance for (B, C, F) was evaluated with unpaired Student's 2-tailed t-test. Statistical significance for the proteomics dataset (D, E) was calculated using a Student's t-test followed by Benjamini-Hochberg FDR correction of 5%. *$p<0.05$, **$p<0.01$, ***$p<0.001$.

### Figure S3

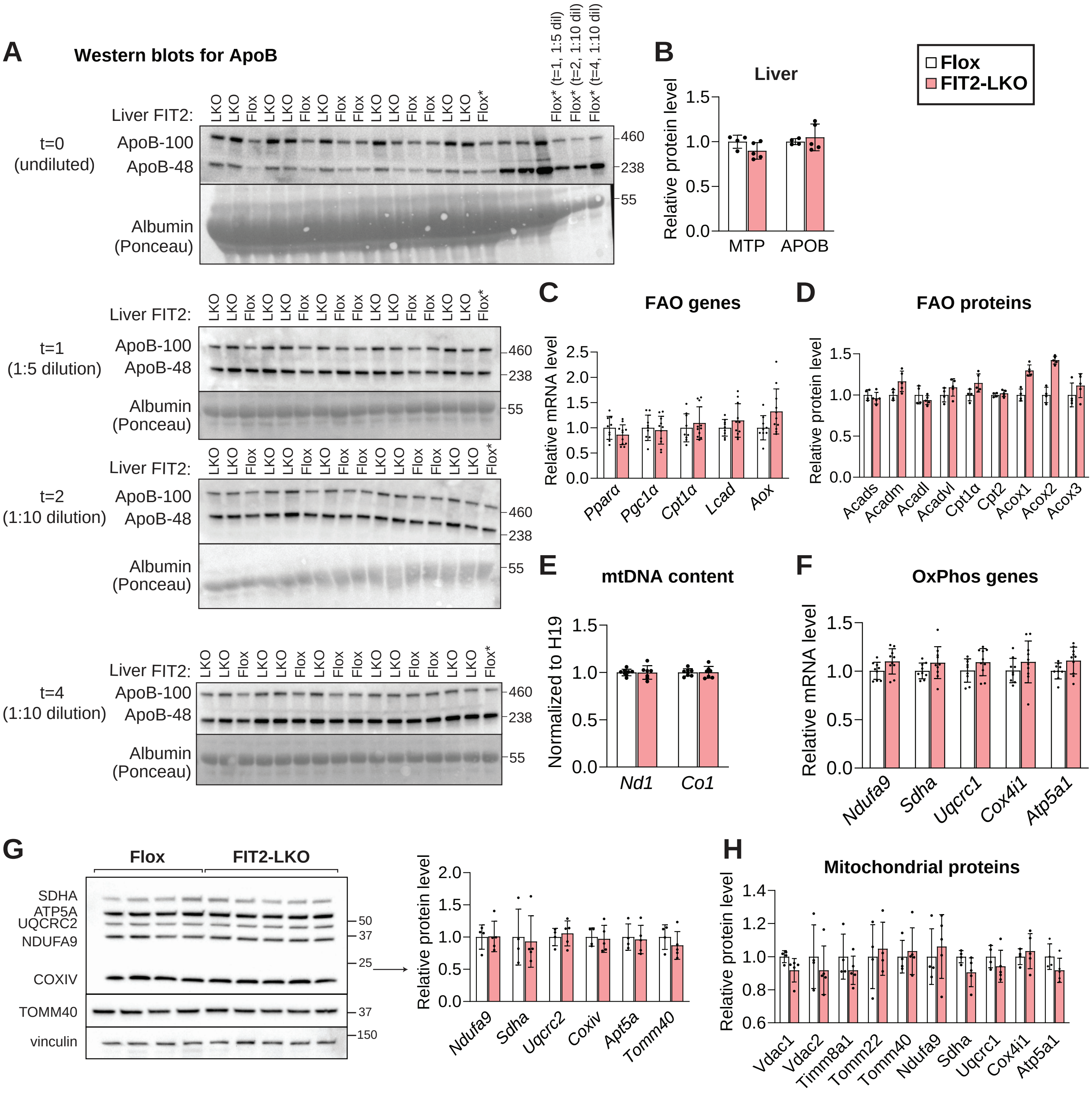


**Figure S3. Corresponding to Figure 3.** (A) Immunoblotting of plasma apoB-100 and apoB-48 used for quantification of ApoB secretion in Figure 5A. (B) Proteomic measurements indicate that amounts of APOB and MTTP protein levels are not altered in the liver between Flox and FIT2-LKO mice (n=4–5/genotype). Genes (C) and protein (D) involved in fatty acid oxidation (FAO) are not reduced in FIT2-LKO livers (n=9–11/genotype). (E) Levels of hepatic mitochondrial (mt) DNA do not differ between genotypes. OxPhos genes (F) and proteins (G, H) are not altered in FIT2-LKO livers. Data represents mean ± SD. Statistical significance for (C, E, F, G) was evaluated with unpaired Student's 2-tailed t-test. Statistical significance for the proteomics dataset (D, H) was calculated using a Student's t-test followed by Benjamini-Hochberg FDR correction of 5%.

**Figure S4.**

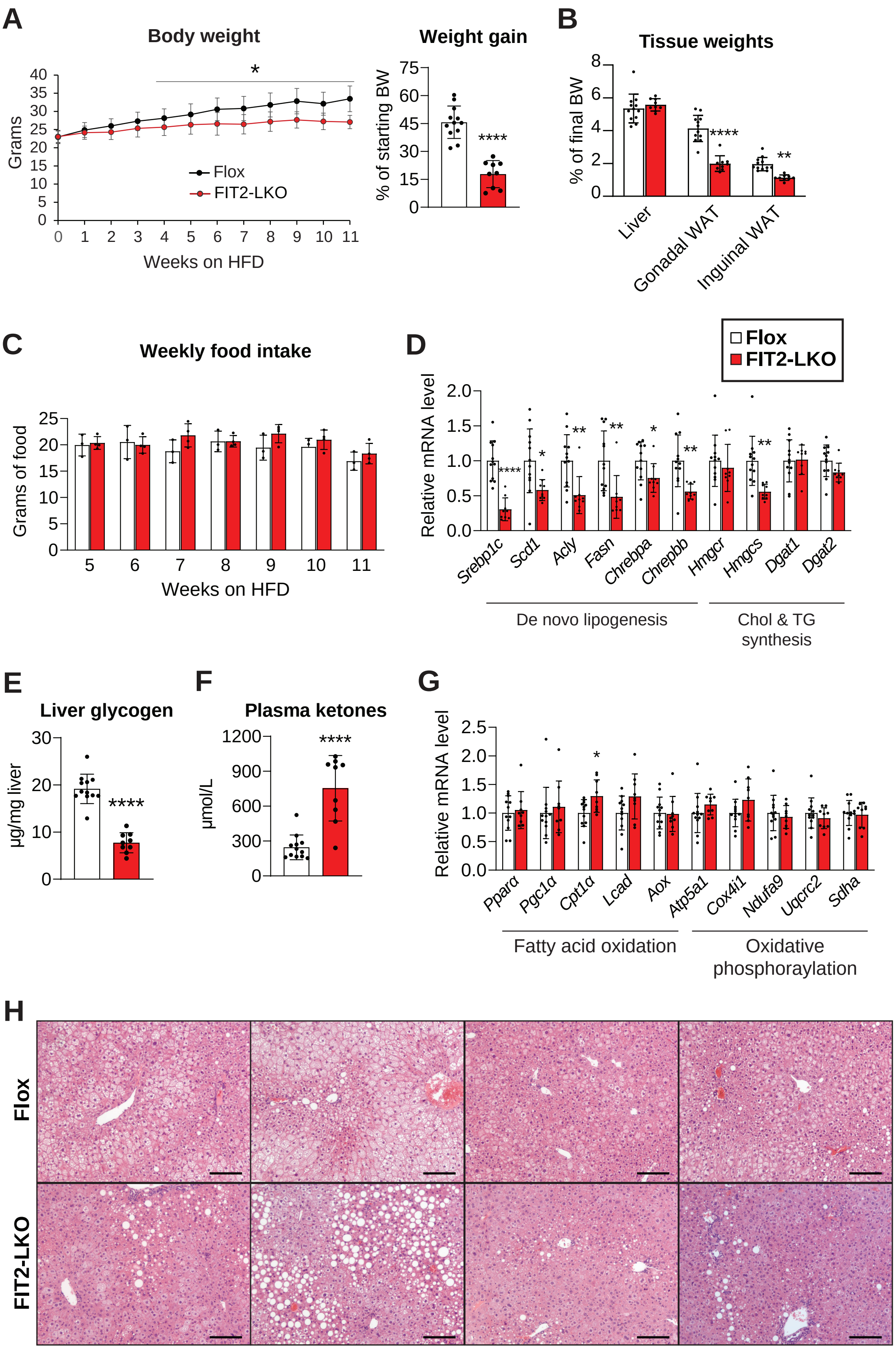
A
Body weight
Grams
Weeks on HFD
Flox
FIT2-LKO
Weight gain
% of starting BW
****
B
Tissue weights
% of final BW
Liver
Gonadal WAT
Inguinal WAT
****
**
C
Weekly food intake
Grams of food
Weeks on HFD
D
Flox
FIT2-LKO
Relative mRNA level
Srebp1c
Scd1
Acly
Fasn
Chrebpa
Chrepbb
Hmgcr
Hmgcs
Dgat1
Dgat2
De novo lipogenesis
Chol & TG synthesis
E
Liver glycogen
μg/mg liver
F
Plasma ketones
μmol/L
G
Relative mRNA level
Pparα
Pgc1α
Cpt1α
Lcad
Aox
Atp5a1
Cox4i1
Ndufa9
Uqcrc2
Sdha
Fatty acid oxidation
Oxidative phosphorylation
H
Flox
FIT2-LKO

**Figure S4. Corresponding to Figure 4.** FIT2-LKO animals weigh significantly less (A) and have reduced adiposity (B) in both gonadal and inguinal white adipose depots compared to Flox mice after an 11-week high-fat diet (HFD) feeding study. (C) Food intake is not different between genotypes. Animals were individually caged and food consumption was measured weekly for 7 weeks (n=3–4/genotype). (D) Under HFD feeding, FIT2-LKO mice exhibit decreased expression of genes related to de novo lipogenesis and cholesterol synthesis. They also show a decrease in liver glycogen content (E) and an increase in plasma ketone bodies (F). The FIT2-LKO mice exhibit little to no change in gene markers for fatty acid oxidation or oxidative phosphorylation (G). (H) Additional images of H&E staining of livers from Flox and FIT2-LKO mice after HFD challenge (n=5–9/genotype). Scale bar=50 µm. Data represents mean ± SD. N=9–12/genotype, unless otherwise noted above for specific experiments. Statistical significance for body weight over time (A, left) was evaluated with repeated-measures ANOVA, and the weight gain (A, right) was evaluated using an unpaired Student's 2-tailed t-test. (B) and (C) were evaluated with 2-way ANOVA with Šidák correction. For (D), statistical significance was evaluated with unpaired Student's 2-tailed t-test for parametric data and a Mann-Whitney U test for nonparametric data. Statistical significance for (E-G) were evaluated with unpaired Student 2-tailed t-test. $^*p<0.05$, $^{**}p<0.01$, $^{***}p<0.001$, $^{****}p<0.0001$.
